# Distinct Dendritic Morphological Changes in the Nucleus Accumbens of Microbiota-deficient Male Mice

**DOI:** 10.1101/2024.02.27.582301

**Authors:** Rubén García-Cabrerizo, Maria Francesca Viola, Pauline Luczynski, Gerard Clarke, John F. Cryan

**Affiliations:** APC Microbiome Ireland, University College Cork, Cork, Ireland; Department of Anatomy and Neuroscience, University College Cork, Ireland; Department of Psychiatry and Neurobehavioral Science, University College Cork, Ireland

**Author notes:** **Correspondence**: John F. Cryan, University College Cork, Cork, Ireland. IUNICS, University of the Balearic Islands and Health Research Institute of the Balearic Islands (IdISBa), Palma, Spain (Rubén García-Cabrerizo). LIMES Institute, University of Bonn, Bonn, Germany (Maria Francesca Viola). Department of Medicine, University of British Columbia, Vancouver, British Columbia, Canada (Pauline Luczynski). These authors contributed equally to the present work.

**Keywords:** Nucleus accumbens, germ-free mice, medium spiny neurons, dendritic spines, gut microbiome

## Abstract

The gut microbiota has been shown to be an important regulator of brain and behaviour. Germ-free rodents are a key model to study the microbiome-gut-brain axis to reveal the microbial underpinnings of diseases, including those related to psychiatric illnesses. The present study evaluated whether the absence of gut microbiota could alter the morphological development of the nucleus accumbens, a brain region located in the ventral striatum involved in stress, mood and addiction. In germ-free mice, there was dendritic hypertrophy of medium spiny neurons in the shell and dendritic elongation in the core. This led to an increase in the number of stubby dendritic spines within the shell and an increase in both stubby and thin spines in the core. Taken together, these results indicate that the gut microbiota is essential for the normal development of the dendritic structure of medium spiny neurons in the nucleus accumbens and that altered remodelling may contribute to maladaptive psychiatric disorders.

## Introduction

One of the most recent exciting advances in biology has been the discovery of how the gut microbiome influences host physiology and behaviour (Cryan & Dinan, 2012; Foster & McVey Neufeld, 2013; Sampson & Mazmanian, 2015; Vuong *et al*., 2017; Sherwin *et al*., 2019; Morais *et al*., 2021). The microbiota-gut-brain axis is a bidirectional pathway through which the brain regulates the activity of the gut and *vice versa*, and is critical for maintaining the homeostasis of the host system (Rhee *et al*., 2009; Collins *et al*., 2012; Fung *et al*., 2017; Cryan *et al*., 2019; Morais *et al*., 2020). Dysregulation of this communication pathway may contribute to the development of diverse psychiatric disorders, indicative of the complex relationship between the gut and brain. A key model system for understanding microbiome-host interactions has been the use of germ-free (GF) mice (mice devoid of all microorganisms since birth), which has provided insight into how the gut microbiome shapes host behaviour, physiology and neurology (De Palma *et al*., 2015; Luczynski, Whelan, *et al*., 2016; Chu *et al*., 2019; Darch *et al*., 2021; Knox *et al*., 2023).

Previous research revealed that GF exhibit differences in central neurochemistry and behaviours compared to conventionally raised rodents (Diaz Heijtz *et al*., 2011; Clarke *et al*., 2013; Spichak *et al*., 2018; Lyte *et al*., 2020). This suggests that the gut microbiota may play a role in the development of neurological and psychiatric disorders by altering the function of the central nervous system. For example, GF mice display changes in behaviour, such as increased stress responses, learning and memory impairments, alterations in social cognition, increased locomotor activity, and attenuation in morphine tolerance, as well as anxiety- and depressive-like behaviours (Gareau *et al*., 2011; Clarke *et al*., 2013; Crumeyrolle-Arias *et al*., 2014; Desbonnet *et al*., 2014; Luczynski *et al*., 2017; Zhang *et al*., 2019). These changes in behaviour may be influenced by the brain reward system, which have been shown to be sensitive to the impact of the gut microbiota (García-Cabrerizo *et al*., 2021).

The nucleus accumbens (Acb) is a key region in the ventral striatum of the mesocorticolimbic pathway that is involved in motivation, reward, learning, motor function and emotional processes (Floresco, 2015; Salgado & Kaplitt, 2015). This structure is comprised of two functionally distinct subdivision called core and shell. The core is associated with goal-directed behaviour, learning, and motivation, whereas the shell integrates reward valence and novelty. Both regions are primarily composed of GABAergic medium spiny neurons (MSNs) expressing dopamine receptors, substance P, enkephalin and other genes (Francis *et al*., 2015). Dysregulation in the Acb has been consistently implicated in stress, mood and addiction, among other disorders, highlighting the critical role of this brain structure in emotional behaviour and as a relevant target for therapeutic exploration. Recent data has implicated different brain areas as targets of microbial signals ranging from hippocampus, amygdala to prefrontal cortex (Cowan *et al*., 2018; Guzzetta *et al*., 2022; Lynch *et al*., 2023; Sharvin *et al*., 2023). However, whether the Acb might be a potential target structure regulated by the microbiota remains unknown. To this end, we investigated whether the absence of gut microbiota affects the normal development of the Acb.

## Material and methods

### Animals

Swiss Webster breeding pairs of GF and CC mice were purchased from Taconic (Germantown, New York, USA) and first-generation male offspring were used in the experiment. Mice were held at the same controlled conditions (temperature 21±1°C, 55-60% relative humidity, 12h light/dark cycle) and fed with autoclaved pelleted diet (Special Diet Services, product code 801010). Mice were divided randomly into two experimental groups in this study: one group for stereology (GF=7, CC=7) and the other one for dendritic morphology (GF=5-6, CC=5-6). Mice were euthanised at 9-10 weeks of age. All experiments were performed in accordance with the guidelines of European Directive 86/609/EEC and the Recommendation 2007/526/65/ EC and were approved by the Animal Experimentation Ethics Committee of University College Cork. Part of the tissues from other brain regions obtained from these animals were previously published in (Luczynski, Whelan, *et al*., 2016; Luczynski *et al*., 2017).

### Stereological Measurement of Nucleus Accumbens Volume

#### Histological preparation

Mice were anaesthetised with sodium pentobarbital and transcardially perfused with 0.1 M phosphate buffer followed by 4% paraformaldehyde. Brains were postfixed in 4% paraformaldehyde for 24 hours, cryoprotected for 48 hours in 30% sucrose solution, flash-frozen in isopentane and stored at -80°C. For each animal, 40 μm coronal sections were cryostat cut, stained with thionin and evaluated under blinded conditions until the final statistical analysis.

#### Analysis of Nucleus Accumbens Volume

The volume of the Acb shell (AcbSh) and core (AcbC) were estimated using the mouse brain atlas (Paxinos and Franklin, 2001). Preliminary studies were used to establish inter-rate reliability with a disagreement of less than 15% on each measurement. Cavalieri’s principle was used to estimate the volume of the Acb regions. The Acb was identified and used as the sample start point at Bregma + 1.97 mm and the end point was chosen at Bregma + 0.61. For each Acb 10-12 evenly-spaced serial sections with a randomised start were studied. Using FIJI software, the regions of interest were manually outlined in each section and converted to area values (mm). The volume was then estimated using the cut section thickness (40μm) and section increment (40μm) combined with the area values.

#### Dendritic Morphology and Spine Density

Mice were euthanised with sodium pentobarbital and transcardially perfused with 0.9% saline. Golgi-Cox staining was performed on the entire brain using a commercially available staining kit (Bioenno Tech, CA, USA) and as previously described (Luczynski, Whelan, *et al*., 2016). All slides were evaluated under blinded conditions until the final statistical analysis.

#### Analysis of dendritic arborization

MSNs were selected for analysis according to the following criteria described (Vyas *et al*., 2002): (1) absence of truncated dendrites, (2) dark and uniform staining, and (3) relatively isolated from neighbouring-stained neurons. For each animal, two neurons in the AcbSh and two neurons in the AcbC per hemisphere were selected for reconstruction. Images were taken through the entire dendritic arbour and 3D reconstructed to calculate dendritic length and branches in Imaris (Bitplane, Switzerland) as previously reported (Luczynski, Whelan, *et al*., 2016). The extent and complexity of dendritic material as a function of radial distance from the soma was performed on 2D images (sholl analysis) of the traced neurons using the plug-in for FIJI. The radius step size was 20μm.

#### Analysis of spine density and total number of spines

Dendritic spines were counted manually using image stacks (100X magnification; N.A: 1.4). For each neuron, 3 segments of approximately 20μm were chosen, based on consistent and dark impregnation throughout the whole dendrite. All protrusions were counted as spines regardless their morphological characteristics if they were continuous with the dendritic shaft. RECONSTRUCT image analysis software was used to classify spines (Risher *et al*., 2014), and were divided into subtypes by measuring the width and length of each individual spine. Spines were classified as filopodia (length value > 2 μm), mushroom (width value > 0.6 μm), thin (length to width ratio > 1), or stubby (length to width ratio ≤ 1). Values were calculated by averaging the spine densities of the sampled segments for each MSN neuron. The total number of spines were estimated by combining the average spine density and the dendritic length for each neuron. The total number of spine subtypes per neuron were also calculated as described previously (Luczynski, Whelan, *et al*., 2016).

#### Statistical analysis

Statistical analyses were conducted using SPSS v.28 (NY, USA). Results are expressed as mean values ± standard error of the mean (SEM). Two-tailed Student’s *t*-test was used to detect group differences (CC vs GF; right vs left hemisphere) in the Acb volume, total dendritic length, number of branches and spine density. For Sholl analysis, a two-way ANOVA followed by a Bonferroni’s post hoc test was used to detect group differences. The level of significance was set at *p* < 0.05. When significance was achieved, percentage changes were calculated using the corresponding control values.

## Results

### Stereological measurement of Nucleus Accumbens volume

14 animals were used in this study (CC: n=7; GF: n=7). Acb volume was not significantly different between hemispheres in either CC or GF mice, therefore hemispheric means were used for all analyses. When compared to CC mice, the Acb of GF mice showed no significant difference in total volume (t_12_=0.42, p=0.68), AcbSh (t_12_=0.26, p=0.80) and AcbC (t_12_=0.47, p=0.65, Figure 1B). There was no significant difference in total brain volume between CC and GF mice (data not shown).

**Figure 1.**
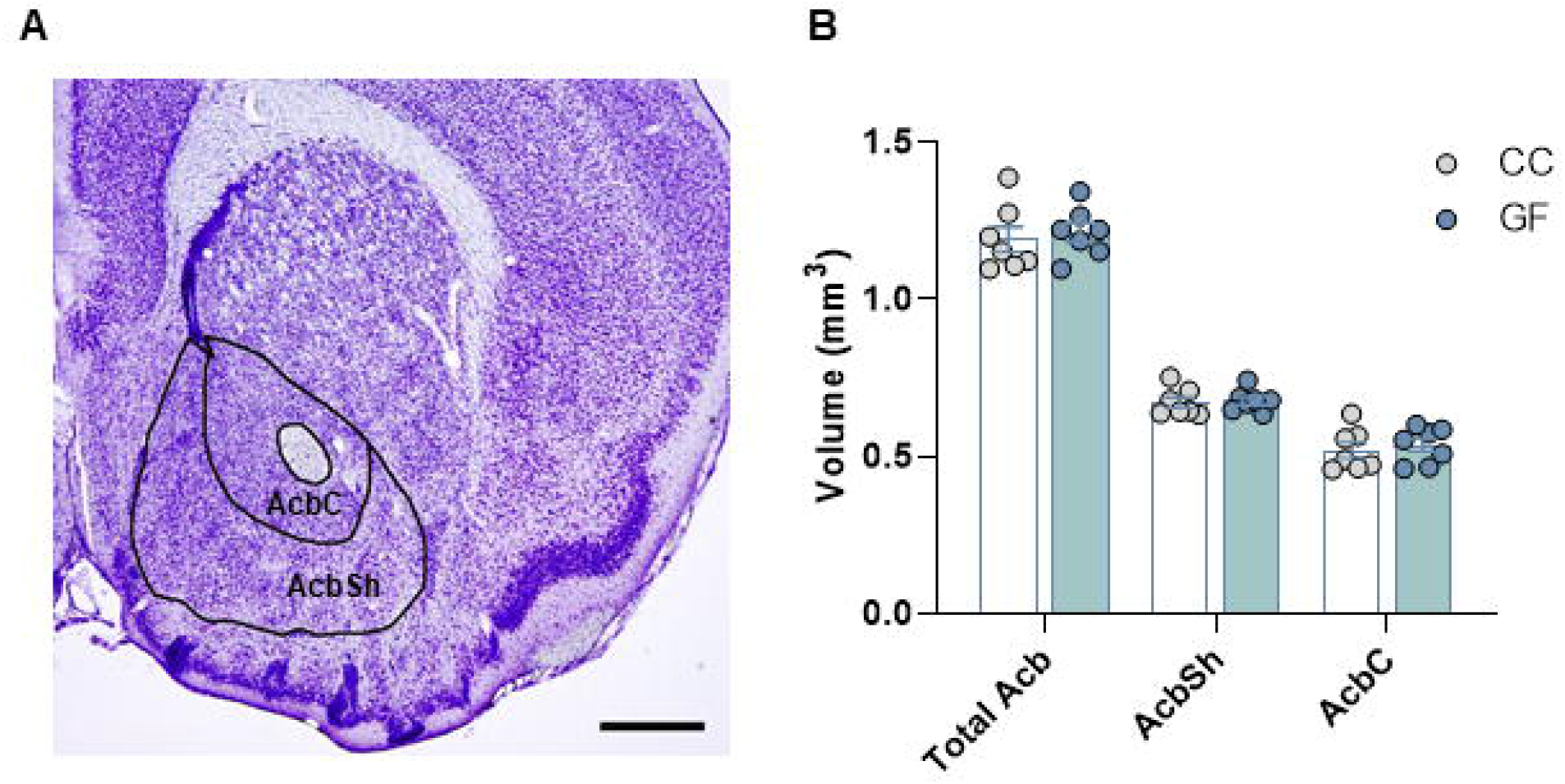
Germ-free status does not change the nucleus accumbens volume compared to CC mice. A) Representative photomicrograph of a thionin-stained section of the Nucleus Accumbens (A; scale bar = 0.5 mm). The volumes of the regions of interest (black lines) were estimated using Cavalieri’s principle. B) Nucleus accumbens shell, core and total volume were not significantly different in GF mice when compared to controls. CC, n=7; GF, n=7. Data represents mean±SEM.

### Dendritic Morphology and Spine Density

In this study, a total of 12 animals (CC: n=5-6; GF: n=5-6) were used for the analysis of the AcbSh and AcbC. For each animal, 4 MSNs per area of interest were analysed for dendritic length and branching (CC: 20-25 MSNs; GF: 20-25 MSNs). Morphometric analysis was performed on the entire dendritic arbor of Golgi-stained MSNs with cell body in either the shell or the core of the Acb.

### Effect of microbiota on the dendritic morphology and spine density in the AcbSh

MSNs in the AcbSh showed significant dendritic hypertrophy in GF mice compared to CC mice (Figure 2A-H). Total dendritic length was increased by 25% (t_8_=5.11, p=0.0009, Figure 2I) and total number of branch points was increased by 22% (t_8_=6.93, p=0.0001, Figure 2J) in GF compared to CC mice. 2D Sholl analysis revealed a significant interaction of group (CC vs GF) and distance from the soma on dendritic density of MSNs in the AcbSh (F_11,88_=4.226, p=0.0001); post-hoc revealed a greatest dendritic extension (35-44%, p<0.05) within the intermediate portions (80-140 μm) of the arbor in GF mice (Figure 2K). The spine density of each MSN was characterised by sampling several segments per neuron. The data set comprised a total of approximately 2500 dendritic spines from 40 neurons. Overall, spine density was not significantly different in GF mice compared to CC mice (Figure 2L); similarly, spine length and width were also unchanged in GF compared to CC mice (Figure 2M-N). There was no significant difference in the density of different spine subtypes in GF animals (Figure 2P), although there was a strong trend toward a loss of filopodia spines in GF mice (t_8_=2.00, p=0.063), however, this effect failed to reach statistical significance.

**Figure 2.**
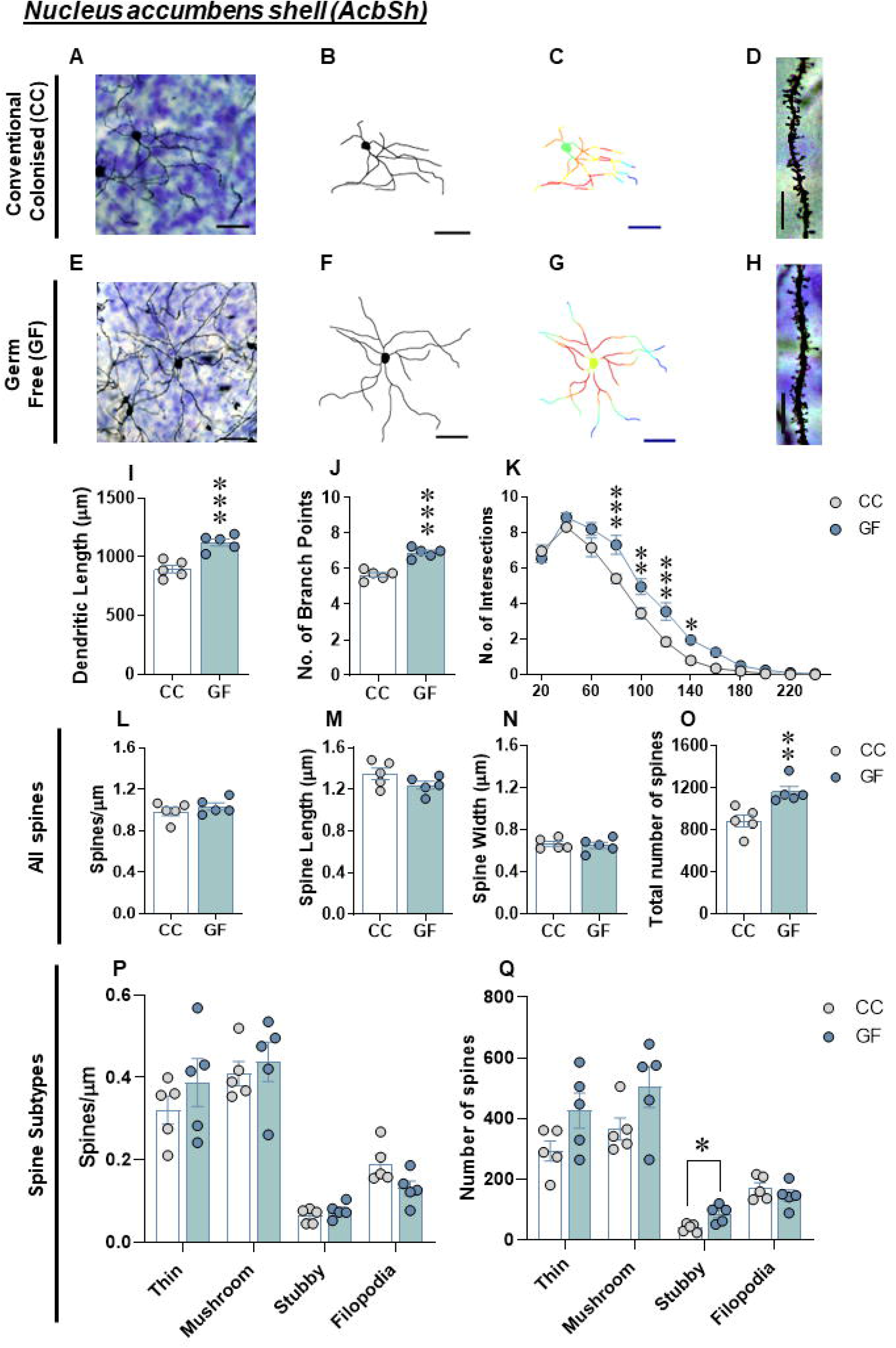
Dendritic hypertrophy of medium spiny neurons in the AcbSh of GF mice. A, E) Representative images of Golgi-stained MSNs in the AcbSh and B, F) resulting computer-assisted morphometric reconstruction. C, G) Sholl analysis was performed on 2D renderings of the neurons (incremental radii from the soma are indicated by the colour gradient). D, H) Representative photomicrograph of AcbSh MSN in CC and GF mice (scale bar = 10 μm). I) Increase in dendritic elongation and J) branching in GF mice when compared to controls. K) 2D Sholl analysis indicates an extension of intermediate dendrites (80-140 μm from the soma) in GF mice. L) There was no difference in spine density, M) spine length and N) spine width of MSNs in the AcbSh in GF mice. O) There was a significant increase in the total number of spines of MSNs in the AcbSh of GF mice. P) There was no difference in abundance of thin, mushroom, stubby and filopodia spines in MSNs of the AcbSh of GF mice when compared to controls. Q) There were no differences in the number of thin, mushroom and filopodia spines, however there was an increase in the total number of stubby spines of MSNs in the AcbSh of GF mice. CC, n=5; GF, n=5. Data represents mean±SEM.

As there was no statistically significant change in spines between groups, we estimated the total amount of spines per neuron by combining spine density and dendritic length. When the dendritic hypertrophy of GF mice was taken into account, there was a 31% increase in the total number of spines of MSNs in the AcbSh of GF mice compared to controls (t_8_=3.616, p=0.007, Figure 2O). In the AcbSh, there was no difference in the total number of filopodia, thin or mushroom spines in GF versus CC mice (Figure 2Q). In contrast, there was a 111% increase in the total number of stubby spines in GF animals (t_8_=3.23, p=0.012, Figure 2Q).

### Effect of the microbiota on dendritic morphology and spine density in the AcbC

MSNs in the AcbC were significantly elongated in GF compared to CC mice (Figure 3A-H). Total dendritic length was increased by 24% in GF (t_10_=2.280, p=0.046, Figure 3I) compared to CC mice, however, branching was not affected by GF status (Figure 3J). 2D Sholl analysis revealed a significant interaction of group (CC vs GF) and distance from the soma on density of MSNs in the AcbC (F_13,130_=2.77, p=0.0016); post-hoc analyses revealed that the greatest dendritic extension (23-41%, p<0.05) occurred within the proximal and intermediate portions (60-100 μm) of the arbour in GF mice (Figure 3K).

**Figure 3.**
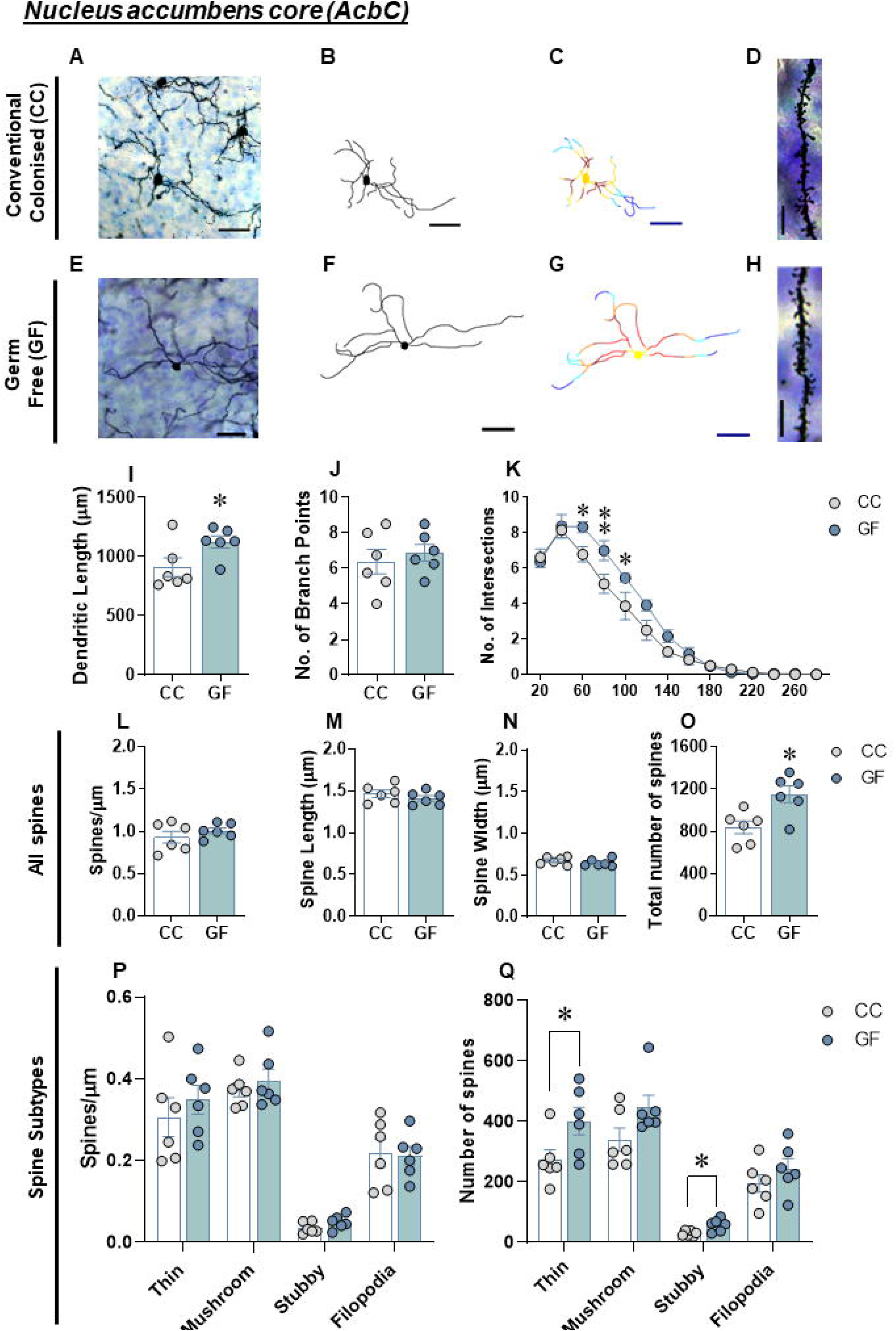
Dendritic elongation of medium spiny neurons in the AcbC of GF mice. A, E) Representative images of Golgi-stained MSNs in the AcbC and B, F) resulting computer-assisted morphometric reconstruction. C, G) Sholl analysis was performed on 2D renderings of the neurons (incremental radii from the soma are indicated by the colour gradient). D, H) Representative photomicrograph of AcbC MSN in CC and GF mice (scale bar = 10 μm). I) Increase in dendritic elongation, J) but without affecting the number of branch points, in GF mice when compared to controls. K) 2D Sholl analysis indicates an extension of proximal and intermediate dendrites (40-100 μm from the soma) in GF mice. L) There was no difference in spine density, M) spine length and N) spine width of MSNs in the AcbC in GF mice. O) There was a significant increase in the total number of spines of MSNs in the AcbC of GF mice. P) There was no difference in abundance of thin, mushroom, stubby and filopodia spines in MSNs of the AcbC of GF mice when compared to controls. Q) There were no differences in the number of mushroom and filopodia spines, however there was an increase in the total number of thin and stubby spines of MSNs in the AcbC of GF mice. CC, n=6; GF, n=6. Data represents mean±SEM.

Multiple segments of each neuron were sampled to assess spine density. Overall, a total approximately 3000 dendritic spines from 48 neurons were counted. Spine density was not significantly altered in GF mice compared to CC mice (Figure 3L). GF status also did not affect spine length or width when compared to controls (Figure 3M-N). The overall density of thin, filopodia, stubby and mushroom spines did not differ in GF compared to CC mice (Figure 3P). When taking the hypertrophic nature of dendrites in GF mice into consideration, there is a 37% increase in the number of spines in the AcbC of GF compared to CC mice (t_10_=3.163, p=0.01, Figure 3O). There was no significant difference in the total number of mushroom and filopodia spines when comparing MSNs in the AcbC of GF to CC mice (Figure 3Q). However, there was a 47% increase in the total number of thin (t_10_=2.278, p=0.046, Figure 3Q) spines and a 91% increase in the total number of stubby (t_10_=3.023, p=0.013, Figure 3Q) spines in MSNs of the AcbC in GF mice compared to controls.

## Discussion

The present study evaluated whether the gut microbiota could alter the morphological development of a key brain region, such as the Acb. Our findings indicate that GF status led to dendritic hypertrophy of MSNs in the shell and dendritic elongation in the core, accompanied by an increase in the number of certain dendritic spines in both regions. Taken together, the results indicate that the gut microbiota plays a significant role in the development of the Acb, and this neuronal remodelling could be involved in the development of certain psychiatric disorders.

Evidence from GF mice have demonstrated an important contribution of gut microbes in shaping brain structures and inducing volumetric alterations of specific brain regions (Luczynski, Whelan, *et al*., 2016; Luczynski *et al*., 2017). However, despite not observing variations in the Acb size or its structural subdivisions in GF mice in the present study, we examined the dendritic morphology and synaptic connectivity of MSNs in both regions of the Acb. Interestingly, dendritic alterations of MSNs in both Acb regions were observed in GF mice. Increases in dendritic length and branch complexity were detected in the AcbSh in GF mice. This dendritic hypertrophy in AcbSh was accompanied by a higher number of stubby spines in MSNs, indicating changes in the distribution of immature spines in GF. In the same way, MSNs in AcbC showed an increase in dendritic elongation and a higher number of stubby and thin spines in GF when compared to CC mice.

Spine alterations in Acb have been reported after chronic stressors and have also been linked with brain reward dysfunctions, demonstrating a common substrate for vulnerability to affective and addictive disorders (Polter & Kauer, 2014). An increase in dendritic elongation of MSNs in the Acb has been linked to stress-induced anhedonia, a core symptom of depression, associated with reductions in motivation (Bessa *et al*., 2013; Francis *et al*., 2017). Furthermore, stress susceptibility promotes the increase of stubby and thin spines in the Acb of mice associated with depressive-related behaviours (Christoffel *et al*., 2011, 2012; Golden *et al*., 2013; Iñiguez *et al*., 2016). Experiments conducted in GF rodents have revealed alterations in stress responses and also changes in depressive-like behaviours, indicating a possible role of the gut microbiota in certain mood disorders (Sudo et al., 2004; Neufeld et al., 2011; Crumeyrolle-Arias et al., 2014; Campos et al., 2016); however, none of these studies have focused on the Acb region. Similar morphological alterations have been described after the exposure to psychostimulant drugs. The repeated administration of cocaine or amphetamine have been linked to increased MSNs firing and dendritic hypertrophy in the Acb, while nicotine administration has been shown to increase dendritic length and spine density in the Acb of rats (Robinson & Kolb, 1999; Brown & Kolb, 2001). This increase in dendritic branching and spine density on MSNs within the Acb could stem the ability of drugs to modify synaptic connections, contributing to the lasting behavioral effects of repeated drug use, including addiction. The dendritic alterations of MSNs observed in the Acb of GF mice in the present work suggests that gut microbes could impact drug reward processes (García-Cabrerizo *et al*., 2021). In fact, gut microbes and certain pathways have been linked with dopaminergic neurocircuitry, reward and motivation (Han *et al*., 2018; Jadhav *et al*., 2018; Dohnalová *et al*., 2022). Changes in the gut microbiome has been also identified in young binge drinkers, pointing out the importance of gut microbes in regulating alcohol craving, social cognition and emotional functioning (Carbia *et al*., 2023). Moreover, negative modulations of the gut microbiota using cocktails of non-absorbable antibiotics alters drug rewarding effects (Kiraly *et al*., 2016; Lee *et al*., 2018; Hofford *et al*., 2021; García-Cabrerizo *et al*., 2023), highlighting the contribution of the gut microbiota to behavioral and molecular responses to drugs of abuse. Recently, marked baseline gene dysregulations have been reported in the Acb of adult germ-free mice compared to CC and microbiome-depleted mice, providing evidence of the potential developmental effects of the gut microbiome on brain signaling and the plasticity response to external stimuli such as drugs of abuse (Sens *et al*., 2023).

These findings should be interpreted with caution, as there are potential secondary influences of the GF mice in relation to the CC group, which might not primarily from their GF status. This could encompass broader alterations in development (i.e., affecting both the brain and body) over multiple generations within a GF colony which could be attributed to epigenetic mechanisms among other variables (Luczynski, McVey Neufeld, *et al*., 2016). Another limitation to consider is that the neuronal populations studied in the Acb are composed of two distinct neuronal types characterized by the expression of different types of receptors (MSNs-D1 and -D2), which respond differently to pharmacological agents and fulfill differential functions despite their similar structural morphology, playing a distinct role both physiologically and in the development of certain psychiatric disorders (Francis & Lobo, 2017). Finally, is worth noting that the scope of the paper is relatively narrow, focusing on one brain region.

Overall, these finding revealed alterations in the morphology of MSNs in the Acb of GF animals. The alterations observed in this key region of the mesocorticolimbic system warrant evaluation of their implications for behavioural and physiological outcomes relevant to stress, depression and addiction. Future investigations should focus on understanding how manipulations of the gut microbiota could alter the fine structure of the Acb and the potential of these novel treatment strategies in certain psychiatric disorders where motivation, reward and emotional disturbances are present. Additionally, including sex as a variable will be crucial, as biological differences between sexes could influence the intricate interplay between the gut microbiota and brain regions, potentially leading to more accurate and nuanced insights (Jaggar *et al*., 2020).

## Abbreviations

(Acb): Nucleus accumbens
(AcbSh): nucleus accumbens shell
(AcbC): nucleus accumbens core
(MSN): medium spiny neuron
(GF): germ-free
(CC): conventionally colonised

## Acknowledgments

R.G-C received funding from European Union’s Horizon 2020 research and innovation programme under the Marie Sklodowska-Curie grant agreement No. 754535. R.G-C is funded by the Spanish Ministry of Universities and co-funded by the University of the Balearic Islands through the Beatriz Galindo program (BG22/00037). APC Microbiome Ireland is a research centre funded by Science Foundation Ireland, through the Irish Government’s National Development Plan (grant No. <u>SFI/12/RC/2273_P2</u>). JFC is funded by the Saks-Kavanaugh Foundation and Swiss National Science Foundation (project CRSII5_186346/NMS2068).

## Authors contributions

RG-C: Data curation, analysis and visualization; writing-original draft; writing-review and editing. MFV: Data curation, analysis and visualization; investigation; methodology writing-original draft; writing-review and editing. PL: Data curation, analysis and visualization; investigation; methodology writing-original draft; writing-review and editing. GC: Resources; visualization; writing-review and editing. JFC: Conceptualization; project administration; supervision; visualization; writing-original draft; writing-review and editing.

